# On the same wavelength: The relationship between neural synchrony and cognitive ability during movie watching in late childhood and early adolescence

**DOI:** 10.1101/2024.08.22.609236

**Authors:** KM Lyons, W Ho, S Saljoughi, AM Owen, RA Stevenson, B Stojanoski

## Abstract

Cognitive development in late childhood and adolescence occurs alongside increasingly complex environments. To examine how cognitive abilities relate to neural processing of naturalistic experiences, we analyzed fMRI data from 309 children and adolescents (ages 7–15; majority White) in the Healthy Brain Biobank. Participants watched a 10-minute clip of Despicable Me during scanning and completed the WISC. Intersubject correlation (ISC) was used to measure brain synchrony during viewing. Adolescents (11–15) with higher cognitive scores showed greater ISC in default mode and frontoparietal networks. In contrast, younger children (7–11) with high scores showed increased synchrony only in somatosensory regions. These findings suggest age- and cognition-related differences in how children and adolescents process complex, real-world stimuli.

---

The period between late childhood and adolescence is marked by tremendous physical, social, and intellectual development. During this time, the brain undergoes widespread structural and functional changes in regions associated with high-level cognition, including the frontal and parietal cortices (Baker et al., 2015; Blakemore, 2012; Giedd et al., 1999; Sowell et al., 1999; Tamnes et al., 2017). These neural changes coincide with improvements in higher-level cognitive abilities, such as working memory, attention, and reasoning (Crone et al., 2004; Gur et al., 2012; Kwon et al., 2002; Luca et al., 2003; Luciana et al., 2005; Theodoraki et al., 2020). Cognitive development is unique for every child, such that these abilities improve at different rates for different children. This individual variation is predictive of many important outcomes, including academic success (Best et al., 2011; Morgan et al., 2019; Rohde & Thompson, 2007; Zaboski et al., 2018), health (Brown & Landgraf, 2010; Reimann et al., 2020; Stautz et al., 2016), and well-being (Nieto et al., 2020; Stern et al., 2017). Coinciding with developing cognitive abilities, the environments children navigate become more complex, with an expanding social circle giving rise to richer and more elaborate experiences (Zelazo, 2004). If cognitive development is associated with more complex experiences, do individual differences in cognitive abilities influence how children experience their world?

Investigating individual differences in how the world is experienced is a challenge, as many of the measures currently available do not reflect how children and adolescents process complex information within the real world. Naturalistic stimuli, such as movies, offer a potentially powerful way to characterize the relationship between individual differences in cognitive development and shared xperiences. Movie watching parallels many real-world experiences; for instance, both require the integration of perceptual and cognitive systems to make sense of complex information that changes across time. When individuals watch the same movie, their brains become highly synchronized (as measured by intersubject correlation, or ISC) with each other (Hasson, Landesman, et al., 2008). Previous work has shown that movies with a narrative lead to greater ISC across the entire brain compared to video clips without a plot (Hasson, Landesman, et al., 2008) and synchrony across participants is absent when they are at rest (Simony et al., 2016). The degree of ISC in adults is predictive of individual differences in emotional processing (Guo et al., 2015), theory of mind (Im et al., 2025), and memory (Baldassano et al., 2017; Chen et al., 2017; Meshulam et al., 2021). Neural synchrony has also been used to assess conscious awareness in clinical populations (Laforge et al., 2020; Naci et al., 2014). These findings have led some to propose that neural synchrony is a measure of shared experiences, particularly when occuring in the frontoparietal and default mode networks (Naci et al., 2014; Nummenmaa et al., 2018; Zadbood et al., 2017).

The frontoparietal network, which is composed of regions in the lateral PFC and intraparietal sulcus, (Cui et al., 2020; Dosenbach et al., 2006; Marek & Dosenbach, 2018; Yeo et al., 2011), is associated with executive processes including working memory, cognitive flexibility, and reasoning (Burzynska et al., 2011; Peters et al., 2016; Satterthwaite et al., 2013; Wendelken et al., 2017), which are essential for plot following. Indeed, synchrony in the frontoparietal network is associated with the degree of suspense and the executive demands of the movie (Naci et al., 2014), and disappears during sedation (Naci et al., 2018). Similarly, the default mode network, which includes regions in the medial prefrontal cortex, posterior cingulate cortex, and inferior parietal cortices (Yeo et al., 2011; Yeshurun et al., 2021), is associated with cognitive processes involving task switching, autobiographical memory, theory of mind, and thinking about the future (Crittenden et al., 2015; Mars et al., 2012; Philippi et al., 2015; Smith et al., 2018; Xu et al., 2016; Yeshurun et al., 2021), all of which are also important for plot following.

Indeed, the frontoparietal default mode networks have been described as the seat of narrative processing (Nguyen et al., 2019). Across several studies, regions within the default mode network show significantly more synchrony when participants are listening to an intact version of a story compared to a scrambled version (Simony et al., 2016), and significant synchrony in frontoparietal and default mode networks was strongly associated with shared or similar interpretation, experience, and memory of the movies (Chen et al., 2017; Nguyen et al., 2019, 2019; Zadbood et al., 2017). Participants who received specific cues to help interpret ambiguous narrative resulted in increased synchrony in many regions of the default mode network, including the medial prefrontal cortex, posterior cingulate cortex, and the temporal parietal junction; this increase in synchrony was not found when participants were not given contextually accurate cues (Ames et al., 2015; Yeshurun et al., 2017). For example, Yeshurun et al., (2017) experimentally manipulated what participants believed about a narrative they were listening to and found this changed the degree of neural synchrony in several brain networks that are associated with social and executive processing, such as the DMN and, inferior frontal gyrus, ventrolateral prefrontal cortex. In fact, similarity in the time course of neural activity in these regions could be used to reliably classify individual based on believed interpretation with an accuracy between 66% to 88% depending on the voxel.

Despite evidence for general changes in neural synchrony at a group level, there is considerable variability in how much people synchronize to each other while processing the same narrative, even when they have not been primed to interpret the story differently (Gruskin et al., 2020; Hasson & Honey, 2012; Lyons et al., 2020; Meshulam et al., 2021; Moraczewski et al., 2018). That is, some individuals show more synchrony, while others show less in response to the exact same stimuli. This suggests differences in synchrony are not due to the properties of the narrative or movie but are influenced by top-down instructions biasing how the plot is interpreted. Why some people in a group synchronize more than others remains an outstanding question (Hasson & Honey, 2012).

That some individuals synchronize to different degrees implies there is a relationship between behaviour and brain responses. However, the degree to which individuals synchronize scales along a continuum of behavioural measures; it is not that those with similar behavioral measures will be similarly synchronized, but individuals on one end of the behavioural spectrum will be more synchronized than individuals on the other end of the spectrum (Chamberlain et al., 2024). One factor underlying variability in degree of synchrony may be the spectrum of cognitive functioning. Indeed, individuals who scored high on measures of social, emotional and cognitive abilities were more synchronized than individuals who scored lower on the same measures (Chamberlain et al., 2024). Given the contributions of various cognitive processes required to follow the plot of a film, differences in cognitive abilities may impact the degree of neural synchrony between individuals watching the same movie. Indeed, this relationship may be stronger during specific stages of development. For instance, as children’s cognitive abilities mature, those with more similar capacities may have more similar experiences of their environment. For example, the degree of synchrony between individual children to a group of adults in response to educational movie clips has been shown to predict academic abilities in each child Cantlon & Li (2013).

However, there are only a few studies examining neural synchrony in children (Cantlon & Li, 2013; Chamberlain et al., 2024; Moraczewski et al., 2018; Richardson et al., 2018) and most have focused on the degree to which a child’s brain synchronizes with a group of adults processing the same narrative, a measure known as neural maturity. Although neural maturity has been found to correlate with brain development, social cognition, and academic scores (Cantlon & Li, 2013; Moraczewski et al., 2018; Richardson, 2019; Richardson et al., 2018), this approach captures a very specific aspect of ISC; the degree to which youth brains functional resemble adult brains. However, this approach does not, intentionally, reflect whether the similarity in brain activity reflects a similar experience. Indeed, adults and children are likely having very different experiences of the movie. For instance, the abilities and preferences often differ qualitatively between children and adults (Carey, 1985; Chandler & Helm, 1984; Goldberg & Thompson-Schill, 2009; Gopnik et al., 2004; Jonauskaite et al., 2019; Luca et al., 2003; Salles et al., 2016), which likely influences how ones brain responds to complex narratives. Second, previous work has shown that children who have more ‘adult-like’ brains, thought to be due to accelerated development, are more likely to have experienced adverse childhood events and tend to have lower cognitive scores (Callaghan & Tottenham, 2016; B. J. Ellis et al., 2022; Shaw et al., 2006; Tooley et al., 2021).

Calculating ISC between similarly aged participants is a more appropriate measure of shared experiences during development (Bacha-Trams et al., 2024). That is, movie watching requires the synthesis of various socio-emotional and cognitive abilities, and the processing and understanding of the plot are shaped by prior biases and preferences. These abilities, biases and preferences are different for children and adolescents, which can lead to very different experiences of the movie. This likely explains why children and adolescents can enjoy the same movie, but for very different reasons. Previous establishing the relationship between degree of neural synchrony and various cognitive and behavioural measures compared children with adolescents and young adults (Chamberlain et al., 2024). However, the degree of neural synchrony is influenced by the group membership (such as family relationship; Bacha-Trams et al., 2024), and computing ISC separately for children and adolescents provides a means of capturing these differences in experiencing the movie and how they might be related to higher-order cognitive abilities.

The current study used a data-driven approach to examine if participants with different scores on the Weschler’s Intelligence Scale for Children (WISC) differ in their degree of intersubject correlations during movie watching. On the basis of the existing literature, it was predicted that participants with higher scores on the WISC would have greater neural synchrony in brain networks associated with plot following, including the frontoparietal network (Naci et al., 2014) and the default mode network (Chen et al., 2017; Nguyen et al., 2019, 2019; Zadbood et al., 2017).

## Methods

### Participants

Data was obtained from the Healthy Brain Biobank (HBN) database (described in detail in Alexander et al., 2017). The HBN is an on-going initiative to collect a large database of children and adolescents, between the ages of 5 to 21. The Chesapeake Institutional Review Board approved the study, and details on the HBN biobank can be found here: http://fcon_1000.projects.nitrc.org/indi/cmi_healthy_brain_network/. Secondary analysis of the HBN data was approved by the institutional research ethics board at Ontario Tech University. For the current study, participants were included if they were between the ages of 7 and 15 and both anatomical and movie fMRI data had been successfully acquired before the year 2020. Everyone included in the current study had written consent obtained from their legal guardians and written assent obtained from the participant. Participants were not excluded based on their handedness or if they had any diagnoses, including neurodevelopmental disorders, to ensure a sample that is representative of the whole population. All participants included in this study had completed the WISC (Weschler, 2014) and were excluded if their full-scale IQ score was less than 70. Participants were also excluded because of failed registration, excessive motion, or if 25% or more of the data contained large ‘spikes’ (significant fluctuations in signal intensity).

Participants included in the final sample (N = 309, see Table 1 for participant demographics) were grouped into two age cohorts (Children and Adolescents) and four IQ bins (Highest, Middle High, Middle Low, and Lowest). Figure 1 displays the distribution of ages and IQ scores. To assess the relationship between ISC and IQ within similarly aged children and adolescents, participants were split (using a median split of their ages) into two equal sized aged groups. Participants in the children group (N = 155) were below the age of 11.2 years, participants in the adolescent group (N = 154) were above the age of 11.2 years. The IQ bins were created using an inter-quartile split of the IQ data, creating a Lowest, Middle Low, Middle High, and Highest IQ group for each age cohort, which ensured equal sample sizes in each group. Based on a chi-square test of independence the groups did not differ in the proportion of male and female participants across each of the groups (X^2^ < 3.01; p > 0.08). Chi-square tests of independence were also run separately for the age cohorts, the groups did not differ significantly in the number of participants with anxiety/mood disorders, autism, or ADHD (all uncorrected p > .30), except for the adolescent Middle High IQ group (N = 1) who had significantly fewer autistic participants compared to the adolescent Lowest IQ group (N = 11, p = .005). The Lowest IQ group in either age cohort did not statistically differ from any other groups in the number of autistic participants (N ranged from 5 to 8). The Lowest IQ groups in both age cohorts had significantly more participants with specific learning disorders (N = 17 in both age cohorts) compared to the three other IQ groups (p < .001, N range from 5 to 9). A one-way ANOVA revealed no differences in mean framewise displacement (FD; Power et al., 2012) between groups based on cognition in children (F = 1.05; *p* = 0.37) and adolescents (F = 0.203; *p* = 0.89). Kolmogorov-Smirnov (KS) revealed no differences in the distribution of mean FD between any groups; smallest KS value was 0.29; and all p-values were larger than 0.08 (not corrected). See supplementary material for more details.

**Figure 1.**
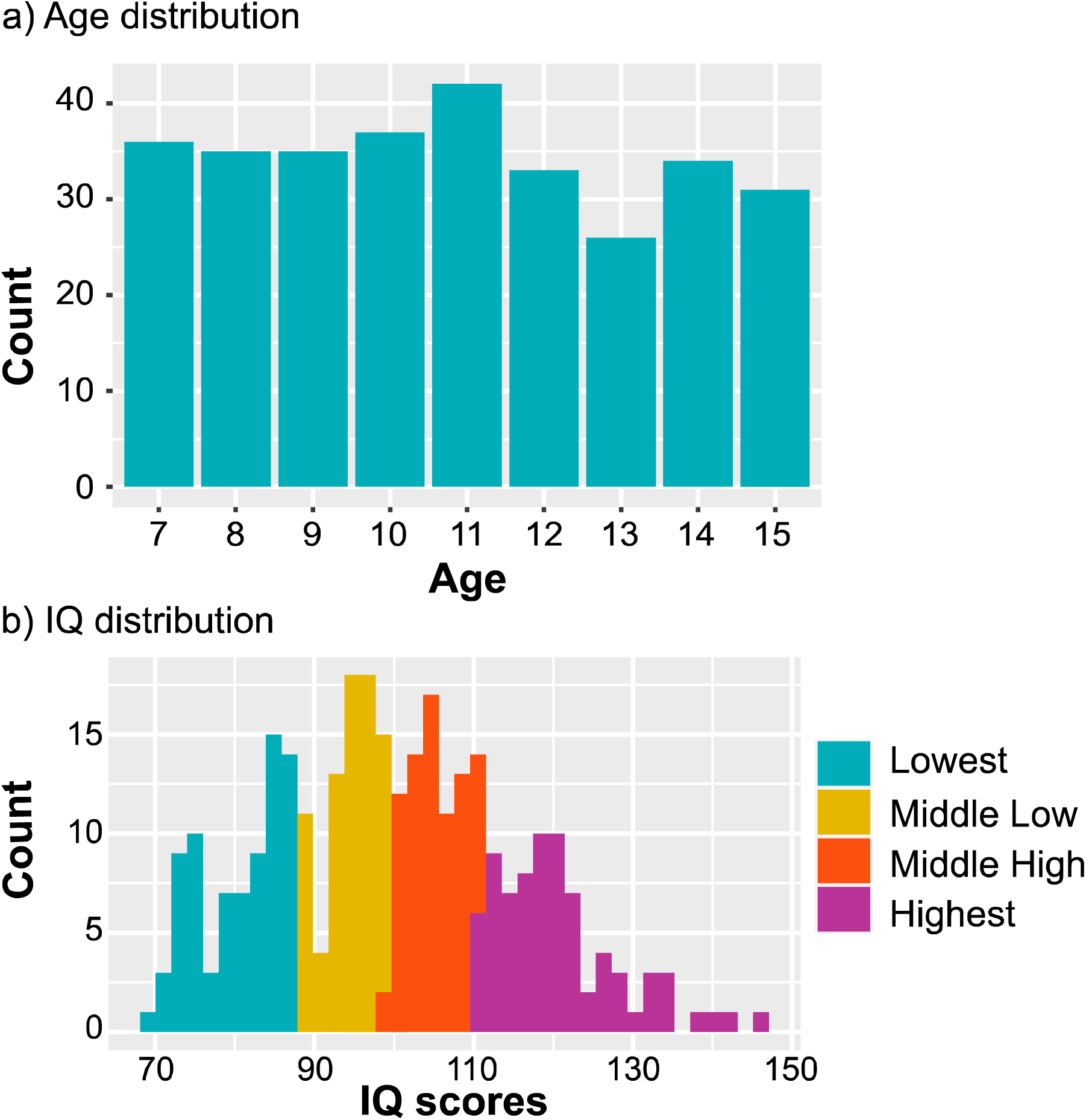
Histograms displaying the distribution of a) age cohorts and b) IQ scores across the four groups.

**Table 1.** Summary of participant demographics.

|  | Group | N | Mean Age(SD) | Mean IQ(SD) | %Female | Motion(SD) | SES(SD) | Diagnosis (N) |  |  |  | %Race |  |  |  |  |  |
| --- | --- | --- | --- | --- | --- | --- | --- | --- | --- | --- | --- | --- | --- | --- | --- | --- | --- |
|  |  |  |  |  |  |  |  | Autism Spectrum Disorder | Attention deficit hyperactivity disorder | Anxiety/mood disorders | Specific learning disorders | White/Caucasian | Black/African American | Hispanic | Asian | Indian | other |
| Children | Lowest IQ | 39 | 9.5 (1.3) | 81.8 (5.8) | 28.20% | 0.28(0.15) | 45.57(15.6) | 6 | 24 | 6 | 17 | 52.80% | 16.70% | 22.20% | 0 | 2.80% | 5.50% |
|  | Middle Low IQ | 39 | 9.2 (1.3) | 96.5 (2.9) | 33.30% | 0.28(0.16) | 48.16(12.67) | 8 | 26 | 9 | 9 | 37.80% | 18.90% | 8.10% | 5.40% | 0 | 29.80% |
|  | Middle High IQ | 39 | 9.2 (1.2) | 106.9 (2.8) | 46.20% | 0.31(0.17) | 52.69(10.27) | 4 | 27 | 11 | 10 | 54.10% | 18.95% | 8.10% | 2.70% | 0 | 16.20% |
|  | Highest IQ | 38 | 8.9 (1.2) | 121.4 (8.0) | 47.40% | 0.29(0.21) | 55.03(9.68) | 4 | 23 | 8 | 5 | 73.70% | 2.60% | 0 | 0 | 0 | 23.70% |
| Adolescents | Lowest IQ | 39 | 13.7 (1.4) | 79.2 (4.5) | 33.30% | 0.25(0.16) | 45.78(15.37) | 11 | 27 | 12 | 17 | 37.80% | 27% | 21.60% | 0 | 0 | 13.60% |
|  | Middle Low IQ | 39 | 13.2 (1.5) | 92.1 (3.6) | 30.80% | 0.22(0.15) | 54.22(12.27) | 5 | 21 | 14 | 9 | 60% | 17.10% | 5.60% | 0 | 0 | 17.30% |
|  | Middle High IQ | 38 | 13.6 (1.3) | 103.1 (3.6) | 34.20% | 0.27(0.19) | 50.62(12.34) | 1 | 19 | 10 | 6 | 50% | 11.10% | 11.10% | 5.60% | 0 | 33.30% |
|  | Highest IQ | 38 | 13.6 (1.3) | 120.5 (8.2) | 34.20% | 0.22(0.13) | 54.56(12.17) | 7 | 21 | 12 | 6 | 64.90% | 8.10% | 5.40% | 0 | 0 | 21.60% |
| Total = |  |  |  |  |  |  |  | 46 | 188 | 82 | 79 |  |  |  |  |  |  |

*Note:* SES scores reflect parent education and income from the Barratt Simplified Measure of Social Status, Motion reflects mean framewise displacement; ASD = autism spectrum disorder and ADHD = attention deficit hyperactivity disorder

### (f)MRI Pre-processing

Functional and structural scans were co-registered and normalized to the Montreal Neurological Institute (MNI) template. We used automatic analysis (AA)(Cusack et al., 2015) and SPM8 to pre-process the functional data. The data were corrected for motion (using six motion parameters: along translational (x,y,z) and rotational (roll, pitch, yaw) axes, spatially smoothed using a Gaussian filter (8 mm kernel), and low-frequency noise (e.g., drift) was removed by high-pass filtering with a threshold of 1/128 Hz. The data were denoised using bandpass filtering, and nuisance regressors that included cerebrospinal fluid, white matter signals, motion parameters, their lag-3 2^nd^-order Volterra expansion (Friston et al., 2000), and "spike" (based on mean signal variance across volumes) regressors (Herzmann et al., 2017; Wild et al., 2017) were used to model sudden intensity (>3 SDs) and motion (mean framewise displacement > 1.3 mm).

### Exploratory whole brain intersubject correlation analysis

The degree of intersubject correlation across the whole brain was calculated using a leave-one-out approach, and this analysis was conducted separately for each group. That is, the pre-processed time course of every voxel was correlated (Pearson and then Fisher z-transformed) between each participant and the average time course of all remaining participants (N-1) from the group across every voxel. A grey matter mask was used to extract r values within grey matter. One-sample t-tests were calculated on the resulting individual brain-wide correlation values. Multiple comparisons were corrected with a false discovery rate (FDR) of 0.05 to generate group maps of significantly correlated voxels.

### Network-based intersubject correlation analysis

The degree of synchronization was computed within seven previously defined functional networks based on the Yeo et al. (2011) parcellation. Hypothesis driven analyses focused on the frontoparietal and default mode networks, given the importance of these two networks for narrative processing. We hypothesized greater neural synchrony in these networks for children and adolescents with better cognitive abilities, reflecting a more similar experience of the movie. However, five additional networks (Visual, Dorsal Attention, Ventral Attention, Somatomotor, Limbic) were included as exploratory analyses. The intra-group ISC for each of these seven networks was calculated using a leave-one-out approach. Specifically, the time course of each network (based on the average time course of each voxel within the network) for each participant was correlated with the average time course of each network for the remaining participants in the group, minus that participant (N-1). Finally, using the rstatix package in R (Kassambara, 2020), a mixed-model ANOVA was conducted to determine if group membership was a significant predictor of intra-group synchronization across the two networks of interest, as well as the five exploratory networks. This was done separately for both age cohorts. The model included ISC values as the outcome variable and group (a between-subject factor) and network (a within-subject factor) as the predictor variables. The networks that showed a significant effect of group were followed up with Welch t-tests (all results were FDR corrected to 0.05).

### Cluster-based intersubject correlation analysis

To ensure the method of grouping participants did not bias the results of the current study, pairwise correlations were calculated in the default mode and frontoparietal networks. This was done by calculating the mean time course (i.e. by averaging across all voxels) separately in both networks for each participant, and then correlating it with every other participant’s mean time course. Finally, a k-means clustering analysis was conducted to determine whether groups of participants could be identified based solely on the degree of neural synchronization, and whether cluster-based group membership was associated with differences in cognitive abilities. The MATLAB function evalclusters was used to identify the optimal number of clusters based on the variance in the data using the Calinski-Harabasz Index, computed over 1000 iterations to minimize the fitting parameter. Based on the groupings generated from this cluster analysis, a single logistic regression analysis with age, full scale IQ, and scores on the five subscales of the WISC as predictors, was computed to investigate which of those factors, if any, (i.e.) significantly predicted the cluster-generated groupings.

## Results

To ensure that potential differences in ISC across the brain are not influenced by motion, we compared mean framewise displacement (FD) between each of the groups. We computed Kolmogorov-Smirnov (KS) tests to determine whether probability distributions differed across the groups. We did not find any differences (across pair-wise group comparisons) in the distribution of mean DF; the smallest KS value was 0.23; and all p-values were larger than 0.22 (not corrected). We also conducted one-way ANOVAs (separately for the child and adolescent cohorts) to determine whether there are differences in the mean of mean FD across groups. We found no differences in mean levels of mean FD across the groups in the child (F_(3,150)_ = 0.73; p = 0.5) or adolescent (F_(3,150)_ = 1.1; p = 0.37) cohorts.

### Whole brain ISC

All age cohorts (Child and Adolescent) and IQ groups (Highest, Middle High, Middle Low, Lowest) showed significant synchrony across the entire cortex (see Figure 2), including much of the visual and auditory cortex, as well as associative regions of the brain.

**Figure 2.**
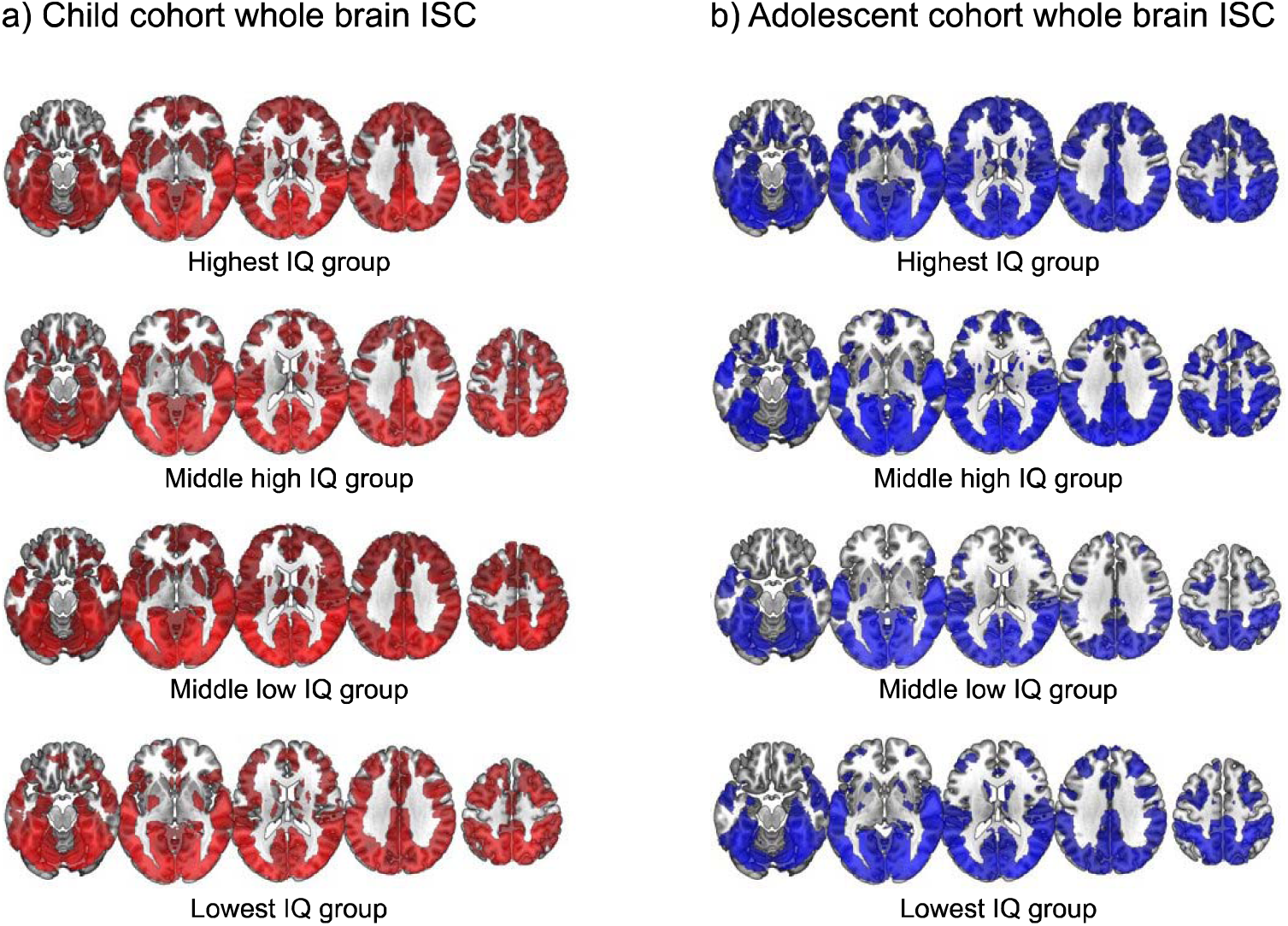
Whole brain spatial maps with significant intra-group ISC displayed for (a) the child and (b) adolescent cohorts. *Note:* All age cohorts and IQ groups had significant synchrony in sensory and associative regions of the brain (FDE corrected p < .05).

### Network based ISC

A mixed effects ANOVA was conducted on both age cohorts separately. ISC values were normally distributed (assessed using Shapiro’s test, p-values ranged from .17 to .72), except in the dorsal attention network for the child cohort (p = .007) and adolescent cohort (p = .04), and in the limbic network for the adolescent cohort (p = .025). There were no significant deviations in the assumption of equal variances in ISC values across groups (assessed using Levene’s test, p-values ranged from .12 to .99). The Greenhouse-Geisser sphericity correction was applied to correct for violations in sphericity. In the child cohort, there was no main effect of group (F_(3, 151)_ = 1.6, p = .19), but there was a main effect of network (F_(4.4, 660.8)_ = 181.3, p < .001) and a significant interaction between group and network (F_(13.1, 660.8)_ = 1.9, p = .02). This significant interaction suggested that the pattern of group differences in ISC values differed across the seven networks. When each network was investigated separately, only the somatomotor network showed a significant group effect (F_(3,151)_ = 5.7, p_adj_ = .007, see Table 3.2). Post-hoc comparisons indicated that children in the Lowest IQ group had significantly less ISC compared to the Middle Low (t_(74.5)_ = 2.94, p = .009), Middle High (t_(69.0)_ = 3.21, p = .006) and Highest IQ group (t_(63.8)_ = 3.90, p = .001).

The adolescent cohort showed a main effect of group (F_(3, 150)_ = 5.0, p = .002), a main effect of network (F_(3.8, 575.9)_ = 144.5, p < .001), but no significant interaction between group and network (F_(11.5, 572.9)_ = 1.3, p = .22). These results suggested that the groups differed significantly in their degree of ISC, but that the pattern of group differences was not significantly different across the seven networks. Post-hoc comparisons revealed that the main effect of group was driven by the Highest IQ group having significantly higher ISC values compared to the Middle High (corrected p value < .001), Middle Low (corrected p value < .001), and Lowest IQ (corrected p value < .001) groups. No other groups were significantly different from each other (uncorrected p values ranged from .221 to .863, corrected p values ranged from .332 to .863; see Table 2).

**Table 2.**
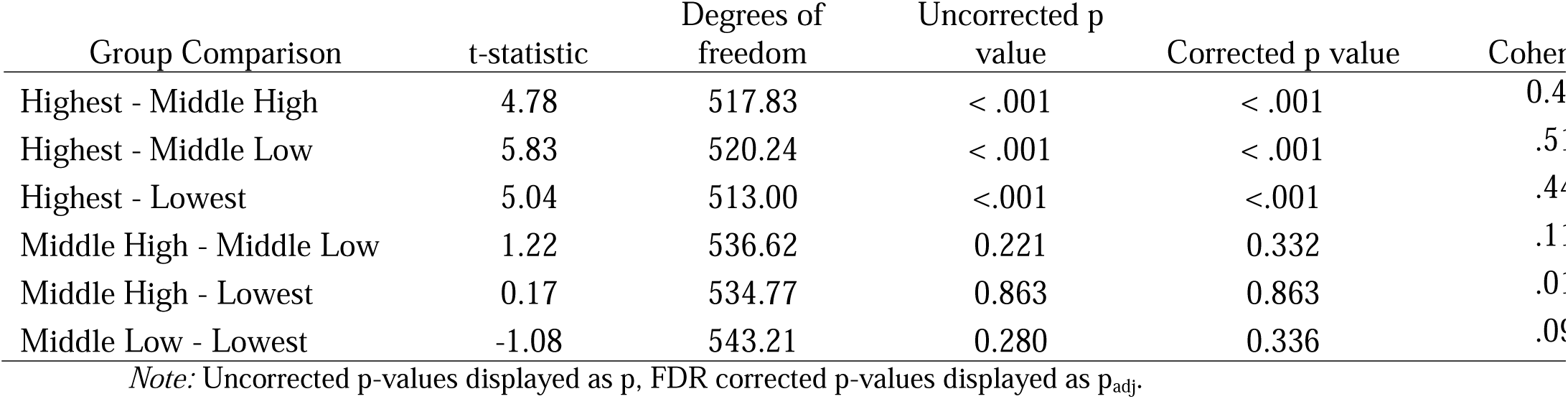
Post-hoc comparisons in neural synchrony across adolescent groups based on cognitive ability.

Despite not finding a significant interaction, we explored the group by network differences to assess our a priori hypotheses that there would be group differences in ISC in the FPN and DMN. When each network was investigated separately, all but the dorsal attention and limbic networks showed a significant group effect (see Table 3 for the statistics for each network). Post-hoc comparisons indicated that for the networks that showed a significant group effect, the Highest IQ group had greater ISC compared to the Lowest and Middle Low IQ groups, although some of these differences failed to reach significance after multiple comparison correction (uncorrected p values ranged from .019 to < .001, corrected p values ranged from .056 to .001, see Figure 3)

**Table 3.**
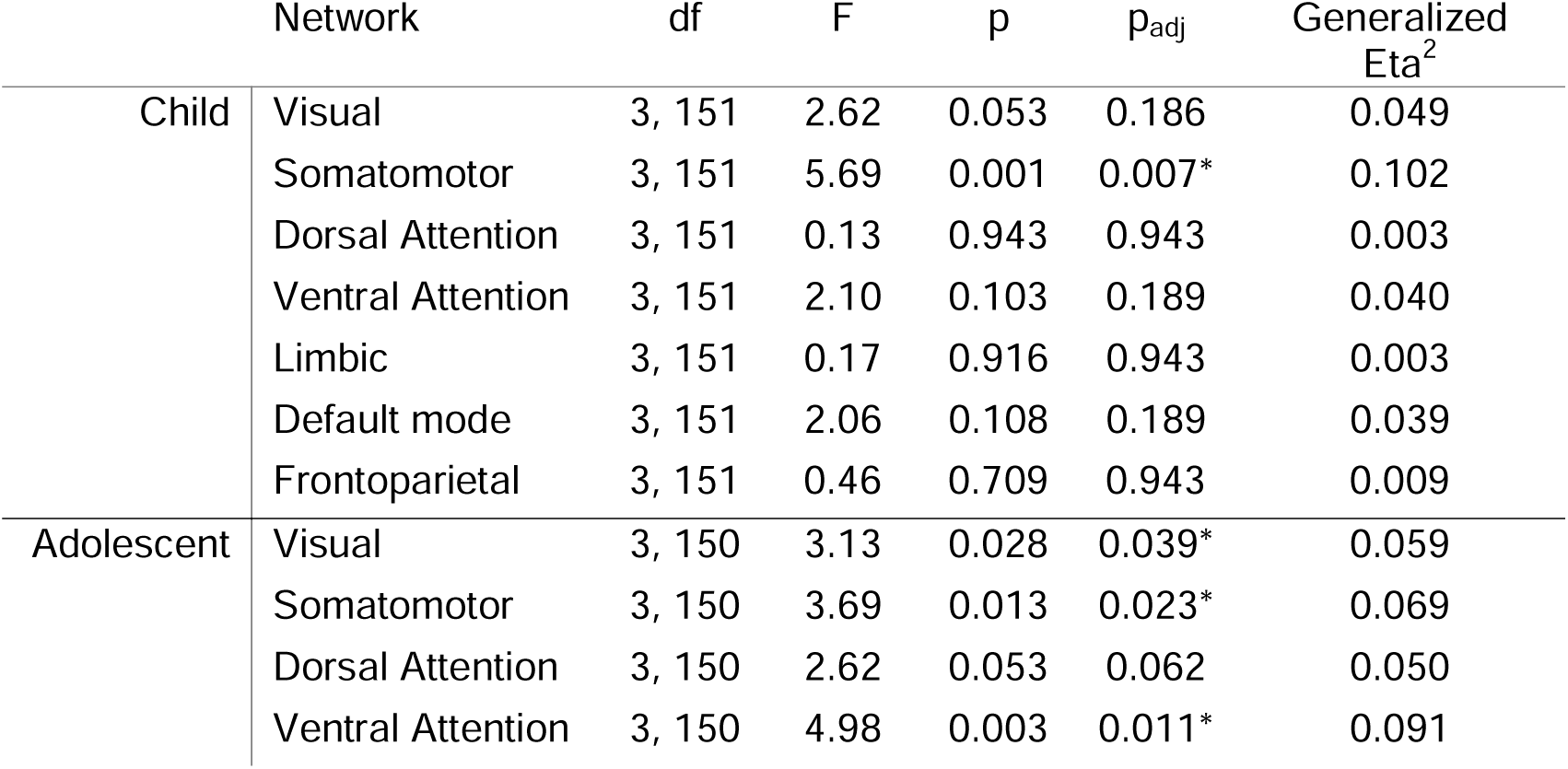
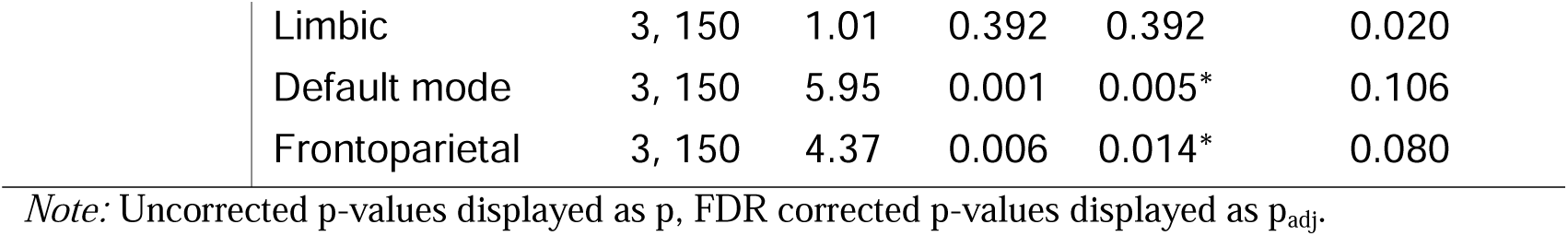
Results from separate ANOVAs calculated to compare ISC in the four IQ groups across the seven networks.

|  | Network | df | F | p | p<sub>adj</sub> | Generalized Eta<sup>2</sup> |
| --- | --- | --- | --- | --- | --- | --- |
| Child | Visual | 3, 151 | 2.62 | 0.053 | 0.186 | 0.049 |
|  | Somatomotor | 3, 151 | 5.69 | 0.001 | 0.007* | 0.102 |
|  | Dorsal Attention | 3, 151 | 0.13 | 0.943 | 0.943 | 0.003 |
|  | Ventral Attention | 3, 151 | 2.10 | 0.103 | 0.189 | 0.040 |
|  | Limbic | 3, 151 | 0.17 | 0.916 | 0.943 | 0.003 |
|  | Default mode | 3, 151 | 2.06 | 0.108 | 0.189 | 0.039 |
|  | Frontoparietal | 3, 151 | 0.46 | 0.709 | 0.943 | 0.009 |
| Adolescent | Visual | 3, 150 | 3.13 | 0.028 | 0.039* | 0.059 |
|  | Somatomotor | 3, 150 | 3.69 | 0.013 | 0.023* | 0.069 |
|  | Dorsal Attention | 3, 150 | 2.62 | 0.053 | 0.062 | 0.050 |
|  | Ventral Attention | 3, 150 | 4.98 | 0.003 | 0.011* | 0.091 |

|  |  |  |  |  |  |
| --- | --- | --- | --- | --- | --- |
| Limbic | 3, 150 | 1.01 | 0.392 | 0.392 | 0.020 |
| Default mode | 3, 150 | 5.95 | 0.001 | 0.005* | 0.106 |
| Frontoparietal | 3, 150 | 4.37 | 0.006 | 0.014* | 0.080 |

*Note:* Uncorrected p-values displayed as p, FDR corrected p-values displayed as p<sub>adj</sub>.

**Figure 3.**
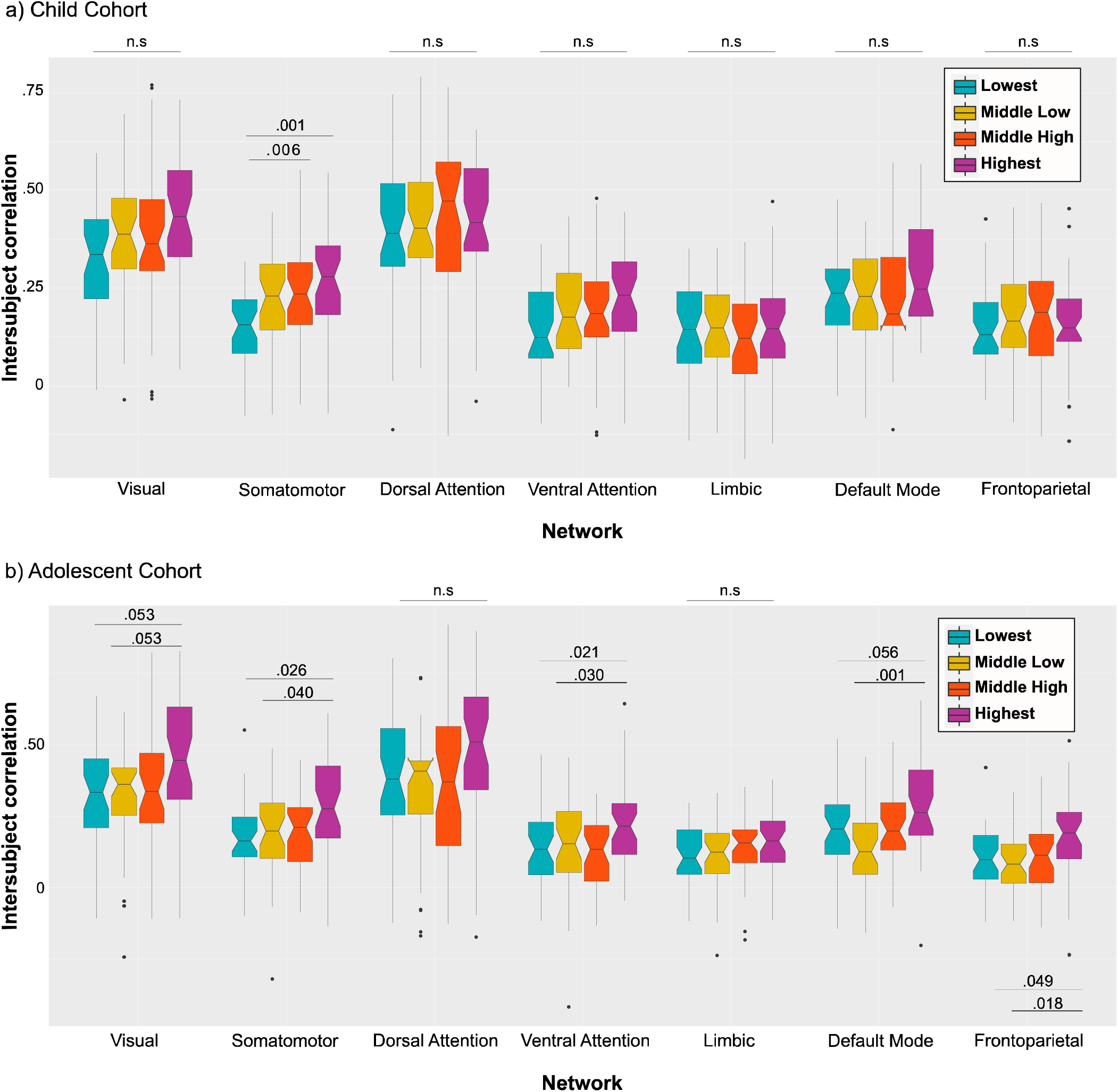
Boxplots displaying ISC for each IQ group across every network for the child cohort (displayed in panel a) and the adolescent cohort (displayed in panel b). *Note:* Reported p-values have been FDR adjusted for multiple comparisons. Middle line represents the median ISC value for each group. The color boxes indicate the interquartile range for each group. Dots indicate outlier ISC values.

### Cluster-based intersubject correlation analysis

To assess the robustness of the intra-group ISC analysis, pairwise intersubject correlations were calculated between each participant and every other participant in the default mode and frontoparietal network (see Figure 4). A k-means clustering analysis indicated that the data best clustered into two groups for both networks: one group with high pairwise correlations and a group with low pairwise correlations (see Table 1 for participant demographics of each group). An exploratory logistic regression analysis indicated that cluster membership in the default mode network was significantly predicted by IQ scores (p = .041), and scores on the visual spatial (p = .025), verbal comprehension (p = .036), and working memory (p = .038) subscales of the WISC. Specifically, those with higher scores were more likely to be part of the high similarity cluster. Age was not a significant predictor of cluster membership in the default mode network. The results of a logistic regression analysis also indicated that cluster membership in the frontoparietal network was significantly predicted by age (p = .006), with participants in the high similarity group being slightly younger than participants in the low similarity group. However, cluster membership in the frontoparietal network was not predicted by full scale IQ or any of the WISC subscale scores.

**Figure 4.**
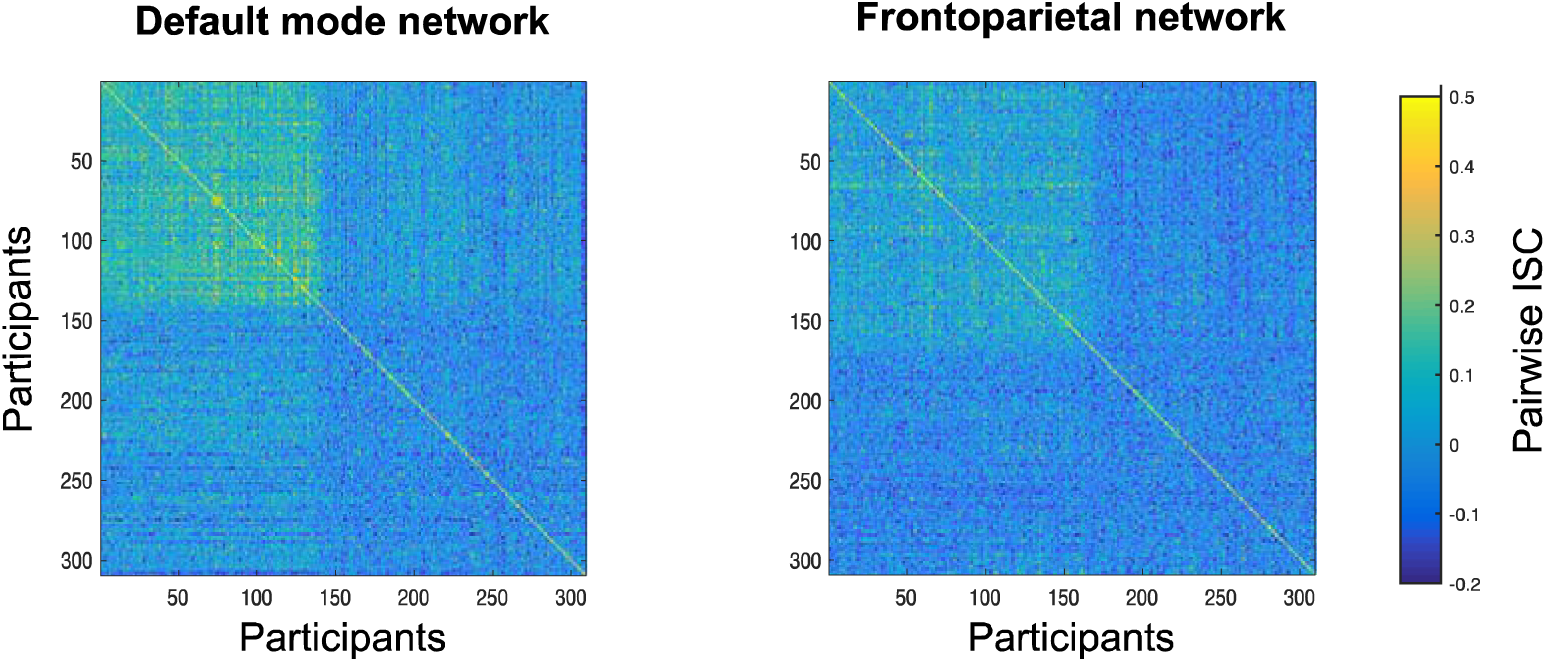
Pairwise correlation matrices across all subjects from all age and IQ groups for the default mode and frontoparietal networks.

**Table 4.** Mean age and WISC scores for the high similarity and low similarity clusters for the default mode and frontoparietal networks.

| Group | *Default mode* |  | *Frontoparietal* |  |
| --- | --- | --- | --- | --- |
|  | High Similarity | Low Similarity | High Similarity | Low Similarity |
| *N* | 138 | 171 | 168 | 141 |
| *Mean Age (SD)* | 11.2 (2.4) | 11.4 (2.6) | 11.0 (2.5) | 11.8 (2.5) |
| *Mean IQ (SD)* | 102 (16) | 98 (15) | 101 (16) | 99 (16) |
| *Mean WMI (SD)* | 100 (14) | 96 (15) | 98 (14) | 98 (16) |
| *Mean VSI (SD)* | 104 (17) | 99 (16) | 102 (17) | 100 (16) |
| *Mean VCI (SD)* | 106 (15) | 102 (15) | 105 (15) | 102 (15) |
| *Mean FRI (SD)* | 103 (15) | 100 (16) | 102 (16) | 100 (15) |
| *Mean PSI (SD)* | 96 (15) | 95 (15) | 96 (15) | 95 (15) |

*Note:* WMI = Working memory index, VSI = Visuospatial index, VCI = Verbal comprehension index, FRI = Fluid reasoning index, PCI = Processing speed index.

## Discussion

The current study found that adolescents (between the ages of 11 and 15) with higher IQ scores showed greater neural synchrony, as measured by intersubject correlation, compared to those with lower IQ scores. Interestingly, of the several of the brain regions that showed this pattern, the frontoparietal and default mode networks specifically have previously been found to be associated with following the plot of a movie. For instance, synchrony in the frontoparietal and default mode networks is modulated based on whether adult participants had more similar (increased) or different (reduced) interpretations (Nguyen et al., 2019), and understanding (Yeshurun et al., 2017) of the narrative of the movie. Additionally, Naci et al. (2014) found that synchrony in the frontoparietal network was associated with the executive load of a movie; that is, time points during movies that are the most engaging were also the periods that show the greatest synchrony in this network. Based on this work in adults, the current study’s findings suggest that adolescents with higher cognitive scores likely experienced the movie in a more similar way compared to those with lower IQs.

However, differences in the degree of neural synchrony between groups of adolescents based on IQ scores were not specific to the default mode and frontoparietal networks. Individuals in the Highest IQ group had significantly greater synchrony in the visual, somatomotor, and ventral attention networks compared to the groups with lower scores. Although synchrony in sensory areas has been shown to occur in the absence of plot following (Hasson et al., 2004; Hasson, Landesman, et al., 2008; Honey et al., 2012; Lerner et al., 2011; Naci et al., 2014), there is evidence that synchrony in sensory areas may be associated with plot following. For instance, several studies have found that individual differences in neural synchrony in the premotor cortex, sensorimotor cortices, and frontal eye fields is predictive of comprehension and memory of the narrative being processed (Meshulam et al., 2021; Yeshurun et al., 2017). These results give rise to an interesting question about individual differences in the relationship between sensory processing and higher-level cognition during movie watching: Do differences in high-level cognition influence lower-level sensory processing during naturalistic stimuli, or do sensory processes form the necessary foundation on which high-level cognitive abilities are built? Addressing this question from a longitudinal perspective will shed light on the neural mechanisms underlying the association between cognitive development and shared experiences.

In contrast to the study’s hypothesis, the pattern observed between adolescents’ IQ scores and neural synchrony in the default mode and frontoparietal networks was not evident in children between the ages of 7 and 11. The only differences in neural synchrony between children in this cohort across groups based on IQ was primarily in the somatomotor network. There are several potential reasons for why the current study failed to find evidence for the hypothesized differences in children. One reason might be that children rely less on high-level cognitive processes when watching movies. However, this explanation seems unlikely because we found significant synchrony across the entire cortex for all four IQ groups in this age cohort. This suggests that they were paying attention to and processing the plot of the movie. Moreover, the movie in the study, *Despicable Me*, is a popular movie for children in this age range, and is in fact part of the highest grossing animated film franchise ever (Jesus, 2017). Another potential explanation for these null results is the asynchronous development of different regions in the brain. It has been proposed that during childhood and adolescence the brain develops along a sensorimotor-association cortical axis, with sensory areas developing before association areas (Sydnor et al., 2021, 2023), and therefore, children are relying on sensorimotor regions to process the movies to compensate for underdeveloped association cortices, including frontoparietal regions (Dong et al., 2021). This does not imply that the functional organization of these networks are not present. Indeed, there is evidence that both sensory (Ellis et al., 2025; Wild et al., 2017) and higher-order networks, such as, theory of mind, pain, and default mode networks develop together (Richardson et al., 2018; Tooley et al., 2021). Although, the underlying architecture of different networks are present early in life, they are in flux, undergoing considerable change (Chen et al., 2022) from early life to early adulthood, and these changes mirror improvements to different cognition abilities (Casey et al., 2000; Dhamala et al., 2021). Relatedly, the parcellation (based on the Yeo et al. (2011) spatial maps) used in this study may not accurately capture functional networks in children. This is particularly relevant for the default mode and frontoparietal networks which undergo considerable change throughout childhood and adolescence, and the adult-based parcellated spatial maps may not completely capture how those networks are parcellated in children (Dong et al., 2021). Consequently, our results may reflect an underrepresentation of the degree to which children’s brains were synchronized during movie-watching. Alternatively, perhaps the current study did not find that IQ was predictive of neural synchrony in children because the method used to group participants may have masked potential differences. The groups were divided arbitrarily based on an inter-quartile/median split, but IQ scores and age are continuous in nature. It is possible that this was not the best way to investigate the relationship between IQ and ISC.

To explore the robustness of the association between IQ scores and neural synchrony, pairwise correlations were calculated across all subjects. This analysis revealed that the data best fit a model with two clusters in both networks: a cluster of high similarity, and a cluster of low similarity. The results of an exploratory logistic regression analysis in the default mode network were consistent with the findings from the IQ grouping analysis in adolescents; participants with more similar time courses were more likely to have higher scores on the WISC (on average, approximately 4 IQ points higher), and on the working memory, verbal, and visuospatial subscales. The two default mode clustering groups did not differ significantly in their age. These results suggest that IQ scores are predictive of synchrony in the default mode network, regardless of the age of participants. Specifically, participants with lower IQ scores, as well as verbal, visuospatial, and working memory abilities, exhibit more idiosyncratic neural responses to a movie compared to those with higher IQs.

Previous work has consistently shown that synchrony in the default mode network is dependent on how someone is engaging with and understanding a movie plot (Ames et al., 2015; Chen et al., 2017; Hasson, Landesman, et al., 2008; Hasson, Yang, et al., 2008; Lerner et al., 2011; Nguyen et al., 2019; Yeshurun et al., 2017). Our findings within the default mode network support the idea that adolescents, and some children who have more matured cognitive abilities, process the movie in a more similar fashion. Higher cognitive abilities may help participants integrate various narrative elements, which may lead to understanding the movie in a more comparable way. It is plausible that having better working memory, verbal, and visuospatial abilities would make following a movie plot less effortful.

The clustering analysis in the frontoparietal network revealed that IQ scores were not predictive of clustering group in this network. These results are in contrast with our other findings that adolescents with the highest IQ group had greater synchrony in the frontoparietal network compared to those with lower scores. One possibility for this conflicting result could be that this effect was subtle in the adolescent group, and therefore was lost when younger participants were included in the analysis.

Conversely, the logistic regression revealed that age was a significant predictor of clustering group in the frontoparietal network. Those in the high similarity cluster were slightly younger, on average by 10 months, than those in the low similarity group. The current sample was skewed towards a slightly younger group (see Figure 1), which may explain why the high similarity cluster in the frontoparietal network contained more younger participants, likely reflecting the idea that similarly aged children experienced the movie. However, this is not the case for all older participants as the high similarity cluster included 15-year-old participants, the oldest age cohort included in this study, suggesting that shared experiences of the movie are not constrained by age. That said, *Despicable Me* was released in the year 2010, so another potential factor contributing to higher synchrony in younger children may be that they were less likely to have seen the movie compared to adolescents, although this information was not available. There is evidence that familiarity with a stimulus leads to a decrease in neural synchrony (Aly et al., 2018; Dmochowski et al., 2012; Sternin et al., 2023). Future studies should investigate how neural synchrony changes across development in a longitudinal design, which would be the best way to disentangle cohort effects from developmental changes.

The current study’s sample was highly heterogeneous; participants were not excluded for their handedness or having a DSM diagnosis, and the range in IQ scores spanned approximately 70 points. Despite this heterogeneity, only two clusters emerged from the pairwise correlations in both networks of interest. Participants in the high similarity cluster had similar patterns of brain activation while watching the movie, whereas those in the low similarity cluster had unique patterns of activation. Although participants in the high similarity cluster in the default mode network were, on average, 4 IQ points higher than the low similarity cluster, the range of IQ scores were nearly identical. Moreover, the age range was also nearly identical (approximately 7 to 15 for both groups). This suggests that in addition to IQ being a significant predictor of neural synchrony in the default mode network, there are other factors that likely predict whether children and adolescents have unique versus similar patterns of brain activation during movie watching. For instance, what explains why a 7-year-old child with an average IQ shows a high degree of similarity in brain activity in the default mode network to a 15-year-old adolescent with a high IQ? Why do two 10-year-old children with similar IQ scores show distinct activation patterns in the default mode network (as is seen in the low similarity cluster)? Future studies should investigate whether these clustering differences are associated with a different interpretation of the naturalistic stimuli. A limitation of the current study is that comprehension data for the movie *Despicable Me* was not collected for the sample used in the current study.

Our study replicates some of the core findings by Chamberlain et al., (2024), and although there are there are several similarities between our findings and those by Chamberlain et al., (2024), there are also important differences. Much like Chamberlain et al., we found that individuals who scored high on different cognitive measures had higher neural synchrony, whereas those who scored lower on the same cognitive scales also had lower degrees of neural synchrony. Unlike Chamberlain et al., who computed ISC and clustered cognitive scores in participants between the ages of 5 and 21, we found find this pattern of results is not consistent across development, and the effect is not consistent across the brain. Our results demonstrate that the link between better cognitive ability and higher neural synchrony was most prominent frontoparietal and default mode networks, but only in adolescents. In children, on the other hand, better cognitive ability and higher neural synchrony was found only in the somatosensory network. Our results, validates those by Chamberlain et al., (2024), but also advances our understanding of neural synchrony and cognition by identifying distinct neural mechanisms underlying this relationship in children and adolescents.

## Conclusion

Overall, the current study found that adolescents with higher IQ scores show greater neural synchrony when watching a movie, compared to those with lower scores. These results suggest adolescents with more mature cognitive abilities may have more similar experiences of naturalistic stimuli. Interestingly, we did not find this pattern in children when participants were grouped by age. In the adolescent group, contrary to our hypotheses, we found differences in neural synchrony appeared most prominently within the default mode network but were less reliably found within the frontoparietal network. This suggests that adolescents with stronger cognitive abilities may be relying on social cognitive abilities, which are supported by several regions of the DMN (Hughes et al., 2019; Li et al., 2014; Spreng et al., 2009), to process the narrative of the move. Once considered task negative, the DMN is important for movie-watching (ref) and other demanding social cognitive tasks such as thinking about oneself (Davey et al., 2016; van Buuren et al., 2010; Wen et al., 2020) or others (Fareri et al., 2020; Li et al., 2014; Schilbach et al., 2008), moral judgement (Chiong et al., 2013; Reniers et al., 2012), and emotional processing (Grimm et al., 2009; Pletzer et al., 2015; Sreenivas et al., 2012). Interestingly, the DMN has also been shown to be linked to other higher-level cognitive abilities (Smallwood et al., 2021), such as, spatial navigation (Spreng et al., 2009), decision making (Marín-Morales et al., 2022; McCormick & Telzer, 2018; Reniers et al., 2012; Smith et al., 2021), and task switching, a key component of the executive functions (Crittenden et al., 2015; Smith et al., 2018).

Future studies should further investigate how cognitive abilities are quantified. Although IQ scores are often treated as a proxy for general cognitive ability, they are also thought to be biased towards certain groups and influenced by socioeconomic factors (Brinch & Galloway, 2012; Capron & Duyme, 1989; Hanscombe et al., 2012; Marks, 2010; Zoref & Williams, 1980). For instance, differences in access to education, home environment, and adverse life events influence a child’s performance on IQ tests (Brinch & Galloway, 2012; Capron & Duyme, 1989; Clearfield & Niman, 2012; Hanscombe et al., 2012; Kira et al., 2012; Ritchie et al., 2013; Ronfani et al., 2015; Saltzman et al., 2006; van Os et al., 2017). In addition, future studies should include objective measures of movie comprehension to link with neural synchrony and cognition. If synchrony in the default mode network is critical for having a similar experience of the movie, including a comprehension measure may explain why some children and adolescents showed a high degree of synchrony in the default mode network, whereas others did not, even among participants with similar ages and cognitive scores.

## Acknowledgements

BS is funded by a Natural Sciences and Engineering Research Council of Canada Discovery grant (RGPIN-2020-05042), and a CIHR Project Grant (487850), and a Canadian Foundation for Innovation John R. Evans Leaders Fund (42163). RAS is funded through two NSERC Discovery Grants (RGPIN-2017-04656 & RGPIN-2024-06233), two SSHRC Insight Grants (435-2017-0936 & 435-2024-1375), a CIHR Project Grant (487850), the University of Western Ontario Faculty Development Research Fund, and a Canadian Foundation for Innovation John R. Evans Leaders Fund (37497), and through a grant from the Canada First Research Excellence Fund (OurBrainsCAN). The funding organizations had no role in the design and conduct of the study; in the collection, analysis, and interpretation of the data; or in the decision to submit the article for publication; or in the preparation, review, or approval of the article. RS, KML, and BS would like to thank Gru for his cunning approach and dedication to completing the project, Dr. Nefario for the constant technical support, and Victor Perkins for continuously challenging us along the way. The authors report no financial interests or potential conflicts of interest.

