## Supplementary Materials for "On the same wavelength: The relationship between neural synchrony and cognitive ability during movie watching in late childhood and early adolescence"

Supplementary Material

Supplementary analysis

**Figure 1**

*Histograms depicting distribution of motion across each the IQ groups for adolescents (older participants) and children (younger participants)*

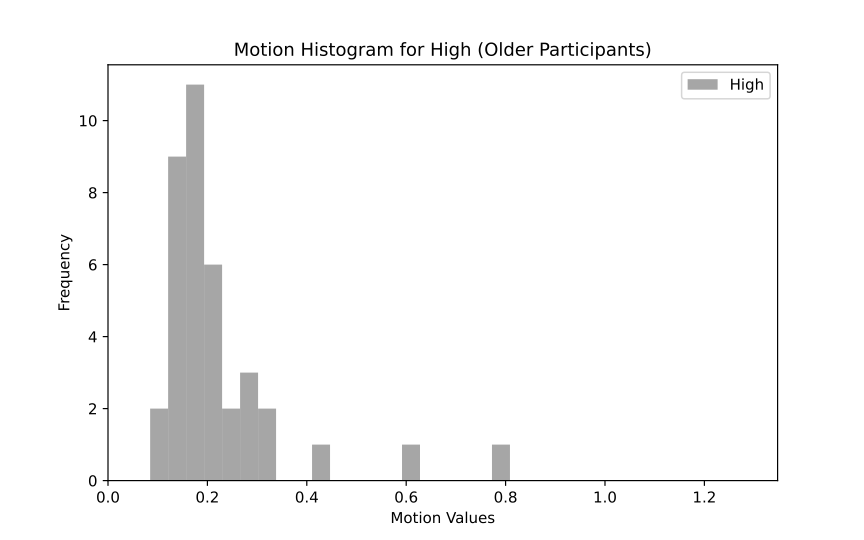

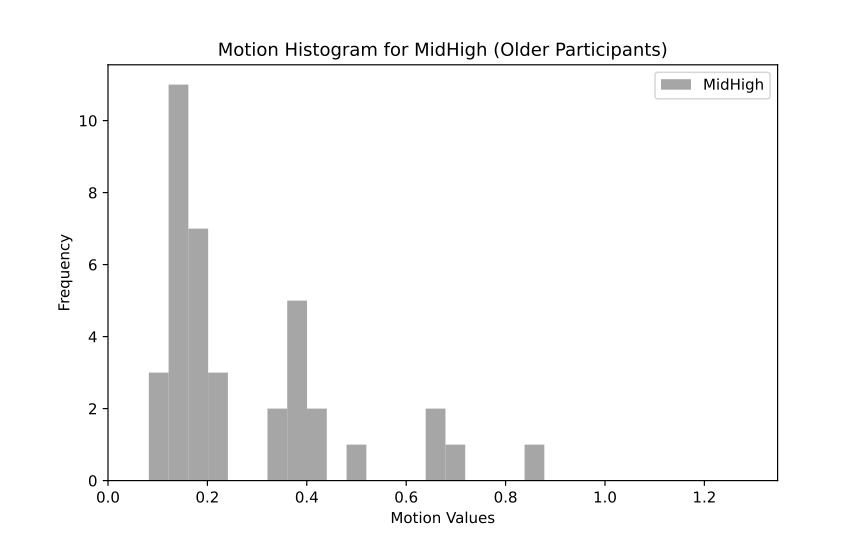

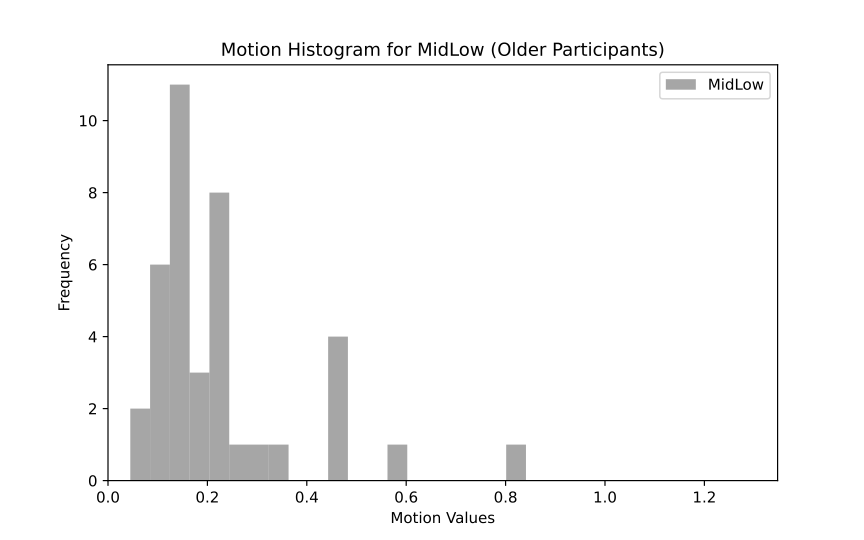

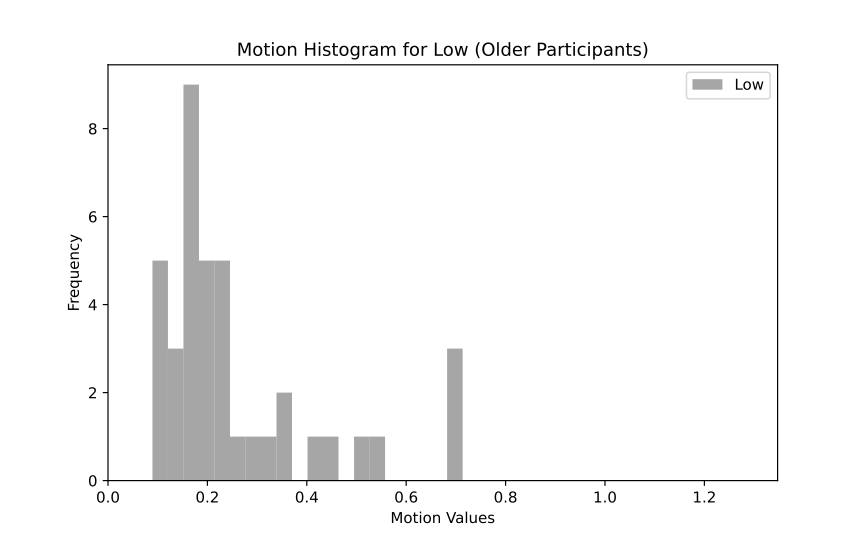

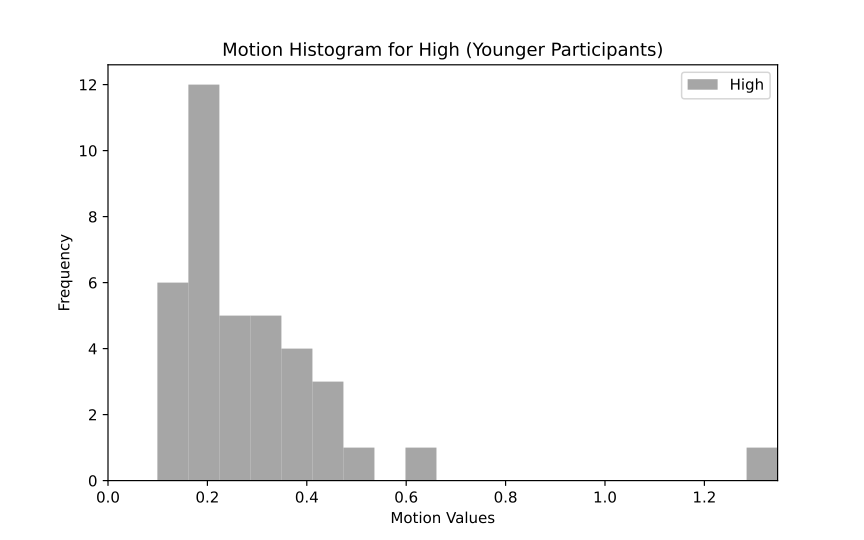

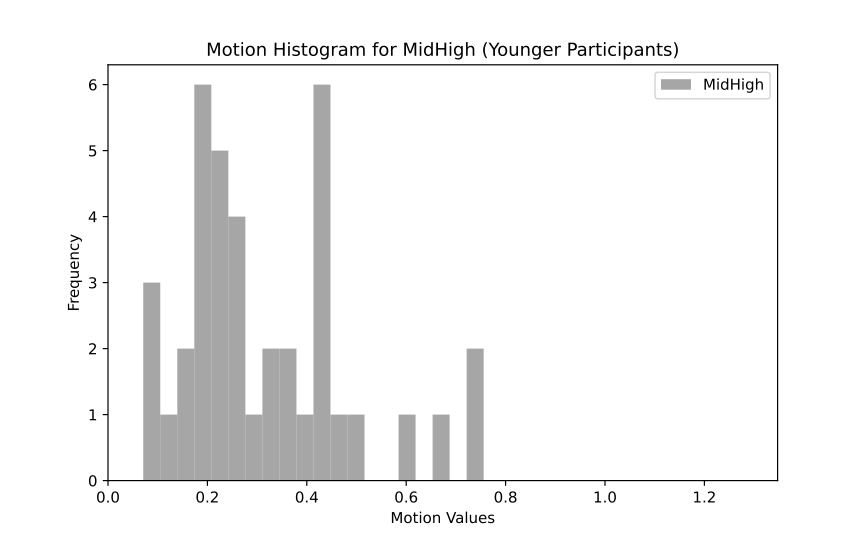

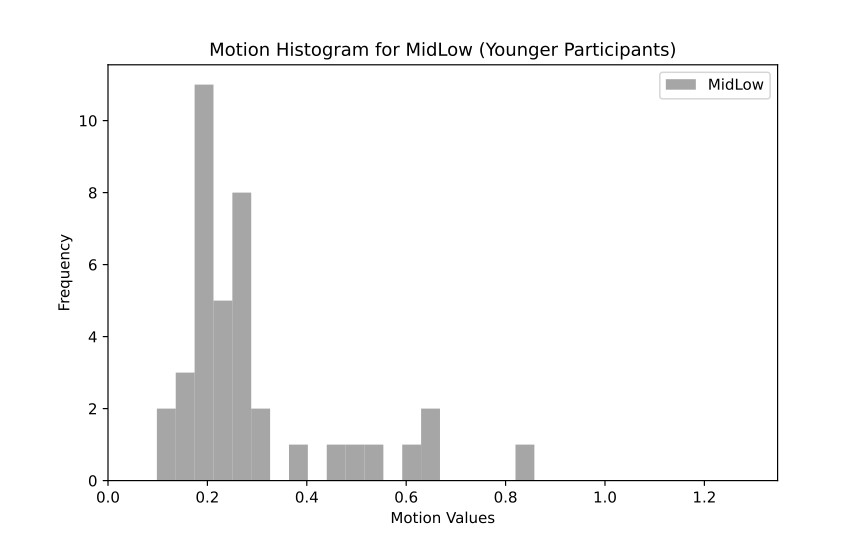

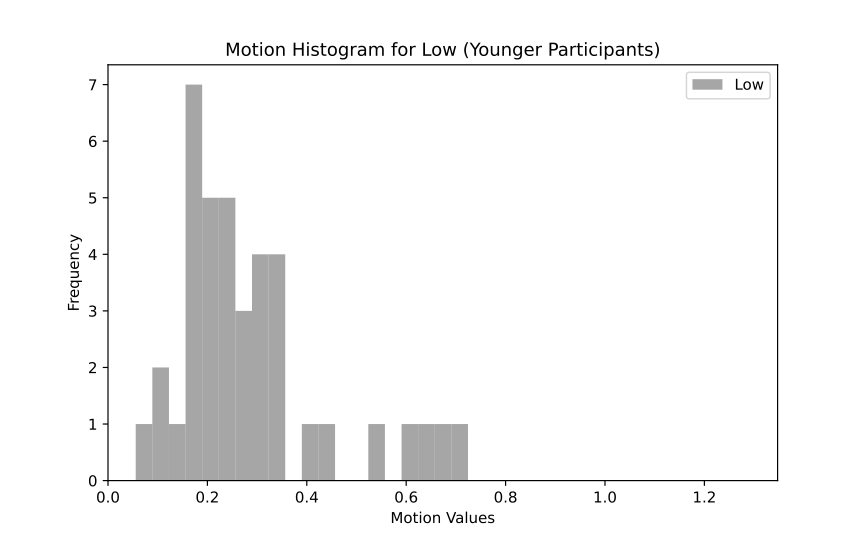

**Supplementary**

*Description of the “Despicable Me” clip*

This clip features a pivotal moment in the narrative where the main character, Gru—a supervillain—adopts three orphaned girls. The clip delves into Gru's internal conflict as he navigates his dual roles: continuing his life as a villian or embracing fatherhood.

The scene begins with the three orphaned girls—Margo, Edith, and Agnes—convincing Gru, their reluctant adoptive father, to read them a bedtime story. This moment highlights the developing emotional bond between Gru and the girls, showcasing themes of attachment and affection.

Following the bedtime story, the narrative progresses to a scene where Gru faces a moral dilemma. He receives a call from his villainous associate, Dr. Nefario, reminding him of his original plan to use the girls for his nefarious schemes. This interaction underscores Gru's internal conflict between his growing affection for the girls and his initial intentions.
